# A phosphoinositide and RAB switch controls early macropinocytosis

**DOI:** 10.1101/2021.11.03.467145

**Authors:** Hélène Spangenberg, Marte Sneeggen, Maria Mateo Tortola, Camila Valenzuela, Yuen-Yan Chang, Harald Stenmark, Camilla Raiborg, Kay Oliver Schink

**Affiliations:** Centre for Cancer Cell Reprogramming, Faculty of Medicine, University of Oslo, Oslo, Norway and Department of Molecular Cell Biology, Institute for Cancer Research, Oslo University Hospital, Oslo, Norway; Interfaculty Institute of Cell Biology, Department of Immunology, University of Tübingen, Auf der Morgenstelle 15, 72076 Tübingen, Germany; Institut Pasteur, Dynamics of Host-Pathogen Interactions Unit and CNRS UMR 3691, Paris France; Division of Molecular and Cellular Biology, National Institute of Child Health and Human Development, NIH, Bethesda Maryland, USA

**Author notes:** Correspondence to: Camilla Raiborg, Kay Oliver Schink.

**Keywords:** micropinocytosis, phosphoinositides, Rab GTPases, endocytosis, membrane transport

## Abstract

Macropinocytosis is a non-selective endocytic process by which cells take up large amounts of extracellular fluids into giant vesicles known as macropinosomes. This mechanism is used by immune cells to sample the surroundings for antigens and can be exploited by cancer cells for nutrient uptake. What determines the fate of macropinosomes after they have been internalized is largely unknown. Here we investigate the role of the phosphatidylinositol 3-kinase VPS34/PIK3C3 and its product phosphatidylinositol 3-phosphate (PtdIns3P) in macropinosome fate determination. Inhibition of VPS34 led to a decrease in macropinosome survival and fluid phase uptake as well as preventing recruitment of early endosomal factors, including the small GTPase RAB5 and its effectors, to the forming macropinosomes. Instead, forming macropinosomes under VPS34 inhibition accumulated regulators of endocytic recycling, including RAB8A, RAB10, RAB11A, and PtdIns4P, which led to fusion of macropinosomes with the plasma membrane.

Whereas RAB5 was critical for macropinosome formation, macropinosome fusion with the plasma membrane depended on RAB8A. Thus, macropinosome maturation is regulated by a PtdIns3P-controlled switch that balances macropinosome fate between the default, endolysosomal maturation and an alternative, secretory route.

## Introduction

Macropinocytosis is an endocytic mechanism that allows cells to take up large amounts of extracellular fluid and substances through the formation of large (>0,5 μm diameter) vesicles [1]. These large vesicles are called macropinosomes and have a variety of functions in different cell types. In immune cells, macropinocytosis is used to sample the surroundings for antigens [2, 3]. Neuronal cells use macropinocytosis to regulate axon growth [4] and to retrieve large amounts of membrane [5, 6].

However, macropinocytosis is also exploited by Ras transformed cancer cells, which frequently upregulate macropinocytosis [7] to support their increased proliferation [8] through uptake of more nutrients, and stimulation of mTORC1 signaling [9]. Likewise, many pathogenic bacteria and viruses stimulate macropinocytosis as a mechanism to enter cells [10–12]. This pathway is therefore an attractive target for therapies against cancer and infections.

In some cell types, macropinocytosis is stimulated by growth factor signaling, while in others it is performed constitutively [13]. In both cases, macropinosome formation requires actin driven ruffling of the plasma membrane. The ruffling plasma membrane forms sheet-like extensions that can close to form cup-like structures. These cup-like structures constrict and are detached from the plasma membrane, forming a macropinosome [1]. The remodeling of the plasma membrane and recruitment of effector proteins required for macropinosome formation are tightly regulated by dephosphorylation of phosphoinositides (PIs). Phosphatidylinositol (3,4,5) trisphosphate (PtdIns(3,4,5)P_3_) is enriched in the preliminary cup-like structures and needs to be sequentially dephosphorylated to PtdIns for successful macropinosome formation [14].

Macropinosomes that successfully undergo this dephosphorylation cascade will have PtdIns on their membranes that can be metabolized by the class III PI3-kinase VPS34 to PtdIns3P [15], the characteristic lipid for early endosomes. At the same time, macropinosomes gain early endocytic factors, like the RAB GTPase RAB5 and its effectors, e.g. Rabankyrin5 and APPL1 [16, 17]. After these early endocytic factors have been recruited, macropinosomes follow a similar maturation route as endosomes, sequentially gaining RAB5, RAB7 and LAMP1 – the characteristic markers for early and late endosomes and lysosomes [18, 19].

One of the key steps during formation of a vesicle is the establishment of correct membrane identity, which determines its fate. Phosphoinositides and RAB GTPases are key determinants of membrane identity. Each compartment has a specific phosphoinositide and RAB code, which can recruit effector proteins and thereby determine the biochemical properties of this compartment [20, 21]. An early-endosomal identity is characterized by the formation of PtdIns3P at the limiting membrane and the recruitment of RAB5 [22]. So far, it is not known how this identity is initially established.

Here, we use macropinosomes as model system to study how vesicles can establish their specific phosphoinositide and RAB code. We show that a VPS34-dependent PtdIns3P pool is critical to recruit RAB5 and to impart endosomal identity to forming macropinosomes. We further show that nascent macropinosomes initially carry secretory markers – PtdIns4P and the GTPases RAB8 and RAB10 – and that VPS34 activity is critical for a RAB- and phosphoinositide switch from a default secretory identity to an endosomal identity.

## Results

### Nascent macropinosomes undergo a VPS34 dependent phosphoinositide switch

In order to visualize macropinosomes, we utilized our recent finding that the PH and FYVE-domain protein Phafin2 labels macropinosomes in two distinct stages – a short, transient localization to nascent macropinosomes directly after scission from the plasma membrane – and a later and sustained localization to the endosomal maturation stages [23]. This dual localization is governed by binding to the phosphoinositides PtdIns4P and PtdIns3P [23].

To probe the establishment of endosomal identity of macropinosomes, we decided to monitor the formation of PtdIns3P – the hallmark lipid of endosomes [24, 25]. We also followed the lipid PtdIns4P [24, 26], which is characteristic for secretory vesicles and the plasma membrane. To detect these phosphoinositide pools, we used genetically encoded lipid probes – mCherry-2xFYVE(HRS) [27] for PtdIns3P and mCherry-2xSidM for PtdIns4P [26], and expressed them together with GFP-tagged Phafin2 as a reference marker. By live cell imaging, we found that newly-formed macropinosomes underwent a phosphoinositide conversion. Nascent macropinosomes initially showed high levels of PtdIns4P, which was rapidly metabolized directly after internalization (Figure 1A). Once PtdIns4P was metabolized, PtdIns3P was generated. It is likely that PtdIns4P is de-phosphorylated to PtdIns, which can then serve as a substrate to generate PtdIns3P, thereby establishing an endocytic identity on the vesicle. The class III PI 3-kinase VPS34 is the main kinase generating PtdIns3P on endosomes [28]. In order to analyze the role of VPS34 and its product PtdIns3P during early steps of macropinosome formation, we used the highly effective and selective VPS34 inhibitor SAR405 [29] to acutely deplete cellular PtdIns3P pools. Treatment of cells with SAR405 blocks de-novo synthesis of PtdIns3P by VPS34. Existing pools are rapidly metabolized, leading to a loss of PtdIns3P from intracellular vesicles within minutes (Figure 1B). When cells expressing Phafin2-GFP and either mCherry-2xFYVE or mCherry-2xSidM were treated with SAR405, we found that localization of mCherry-2xSidM was not affected; rather, this probe showed a slightly longer localization to macropinosome membranes (Figure 1C). In contrast, localization of mCherry-2xFYVE was completely lost, as expected (Figure 1C). Phafin2 only localized to nascent macropinosomes, consistent with earlier findings that its localization to mature macropinosomes is dependent on VPS34-generated PtdIns3P [23]. We also found that this localization was slightly longer than in non-treated cells, similar to the localization of mCherry-2xSidM. We conclude that nascent macropinosomes undergo a switch from PtdIns4P to PtdIns3P as they acquire endosomal identity, and that this depends on VPS34.

**Figure 1.**
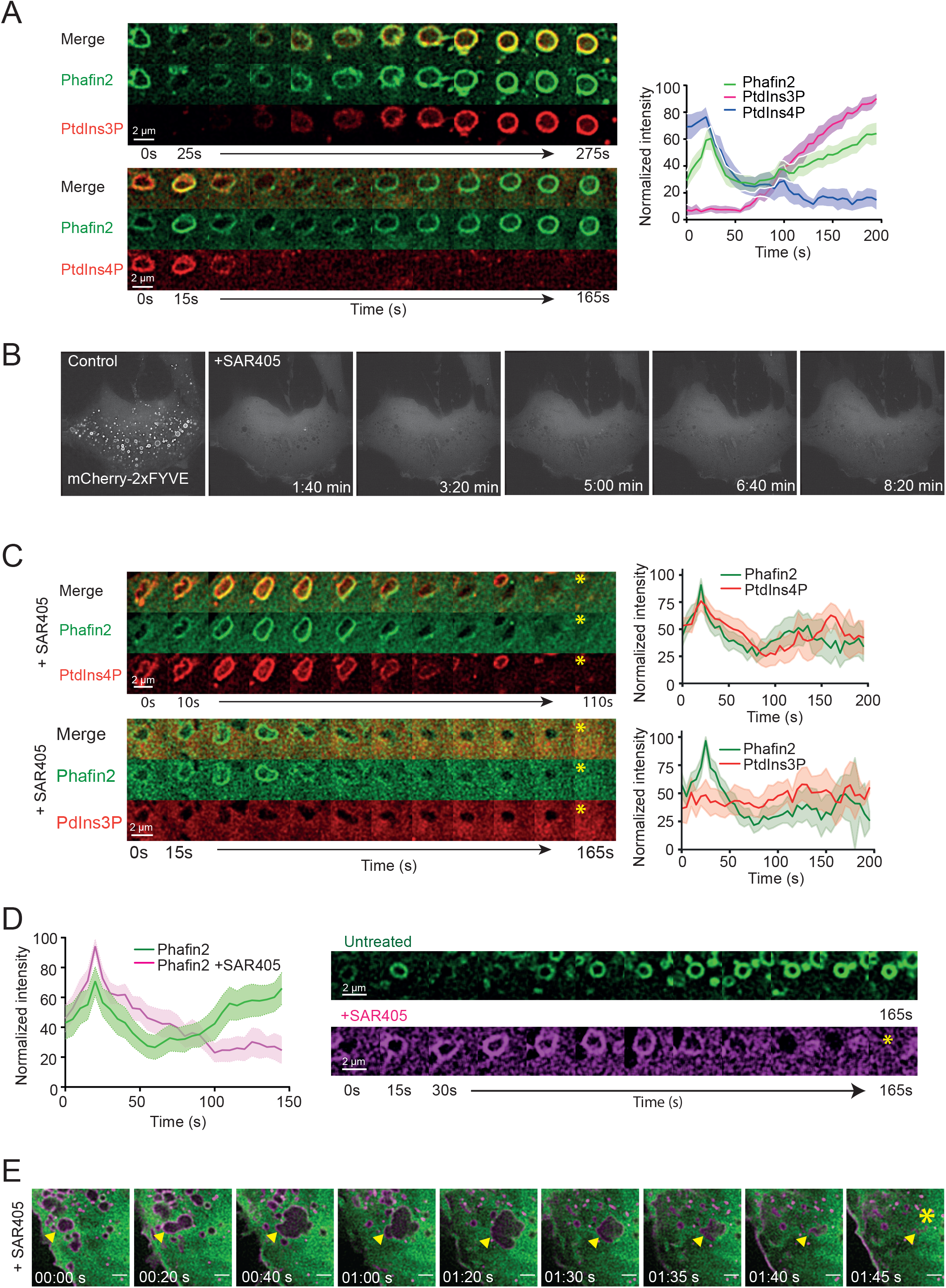
Newly formed macropinosomes undergo a VPS34 dependent phosphoinositide switch (A) hTert-RPE1 cells stably expressing the macropinosome marker Phafin2 were transiently transfected with the PtdIns3P probe 2xFYVE or the PtdIns4P probe 2xSidM and analyzed by live cell imaging. Cells were imaged every 5 seconds. Phafin2 shows a dual localization to forming macropinosomes – one transient early one and a second, later and more stable one. During the first Phafin2-peak also the PtdIns4P probe 2xSidM is recruited to macropinosomes, while the PtdIns3P probe 2xFYVE localizes to macropinosomes during the second, more stable Phafin2 recruitment (mCherry-2xFYVE n= 25; mCherry-2xSidM n=31; mean + 95% CI). Representative sequential images of the recruitment of the lipid probes to the Phafin2-positive macropinosomes are shown. (B) Sequential images of live cell imaging with the PtdIns3P probe 2xFYVE. Upon inhibition of VPS34 with SAR405 (3 μM) the mCherry-2xFYVE probe is no longer on endosomal structures, but in the cytosol. (C) Upon VPS34-inhibition PtdIns4P is stabilized on macropinosomes, while PtdIns3P is lost. Using 2xSidM probe to detect PtdIns4P and 2xFYVE probe to detect PtdIns3P, we examined the effect of Vps34-inhibition on both lipids on macropinosomes. PtdIns4P (n=30; mean +95% CI) seemed to be stabilized for a longer time on newly formed macropinosomes, while PtdIns3P (n=22, mean +95% CI) generation was completely inhibited. Representative sequential images of mCherry-2xSidM and mCherry-2xFYVE recruitment to forming macropinosomes (Phafin2-GFP, mCherry-2xSidM and Phafin2-GFP, mCherry-2xFYVE) are shown. Asterisk indicates end of macropinosome lifetime. (D) Live imaging showing that the recruitment of the macropinosome marker Phafin2 is affected by inhibition of VPS34. Under untreated conditions, Phafin2-GFP shows its biphasic localization to forming macropinosomes. Upon VPS34-inhibition (3 μM SAR405) only the first Phafin2-GFP recruitment is observed, while the second Phafin2-GFP recruitment to macropinosomes is abolished (untreated n= 30 macropinosomes in 7 cells; SAR405 treated n=36 macropinosomes in 5 cells, mean + 95% CI). Representative sequential images are shown of Phafin2-GFP recruitment to the forming macropinosomes in the absence or presence of the VPS34-inhibitor SAR405. (E) Live imaging of hTert-RPE1 cells stably expressing mCherry-MyrPalm (magenta) and mNeonGreen-2xFYVE (green) in the presence of VPS34-inhibitor SAR405 (3 μM). The montage shows the formation of a macropinosome as indicated by a yellow arrow and follows its progress after internalization. The disappearance of the macropinosome is marked by an asterisk. Scale bar 2 μm.

### PtdIns3P depleted macropinosomes re-fuse with the plasma membrane

While tracking macropinosomes in cells treated with VPS34 inhibitor SAR405, we noticed that these vesicles invariably disappeared shortly after their formation (Figure 1D (asterisk)). Newly-formed macropinosomes still gained the first Phafin2 localization – suggesting that they successfully formed and abscised from the membrane [23], but then rapidly disappeared. High resolution imaging of SAR405 treated cells expressing the membrane marker MyrPalm-mCherry and mNeonGreen-2xFYVE showed that these cells could still form macropinosomes – as judged by the appearance of large vesicles – which then shrunk and disappeared (Figure 1E, movie S1).

In order to understand this finding, we used live cell imaging to follow the fate of Phafin2-lablled forming macropinosomes with and without VPS34-inhibitor SAR405 in a cell line stably expressing Phafin2-GFP. In untreated cells, ~65% of all newly-formed macropinosomes matured to the endosomal stage (Figure 2A). In SAR405-treated cells, we found that macropinosomes showed two distinct fates, depending on their maturation state. Macropinosomes forming after the addition of SAR405 invariably collapsed and disappeared (Figure 2A), whereas preexisting RAB5 positive macropinosomes were largely unaffected by SAR405 (Figure 2B). We also compared the lifetimes of newly forming macropinosomes in untreated and in VPS34-inhibited conditions and observed a reduction from a mean lifetime of 465s in control cells to 135s in SAR405-treated cells (Figure 2C). Since the survival of nascent macropinosomes was significantly decreased upon VPS34 inhibition, we investigated whether this was also reflected in fluid phase uptake by macropinocytosis. To that end we incubated cells with fluorescently labelled 70 kDa dextran and measured dextran uptake by flow cytometry. Cells treated with the macropinocytosis inhibitor EIPA [30, 31] or with SAR405 showed a strong reduction in fluid phase uptake of dextran compared to untreated cells (Figure 2D). To investigate whether this reduction of fluid phase uptake upon SAR405 treatment is caused by a decreased rate of macropinocytosis, we performed live cell imaging and quantified the number of forming macropinosomes in untreated, DMSO- and SAR405-treated cells using the first, VPS34-independent recruitment of Phafin2-GFP as a marker. The number of forming Phafin2 positive macropinosomes was unaffected in all conditions (Figure 2E), indicating that the decrease in fluid phase uptake was not due to a defect in macropinosome initiation.

**Figure 2.**
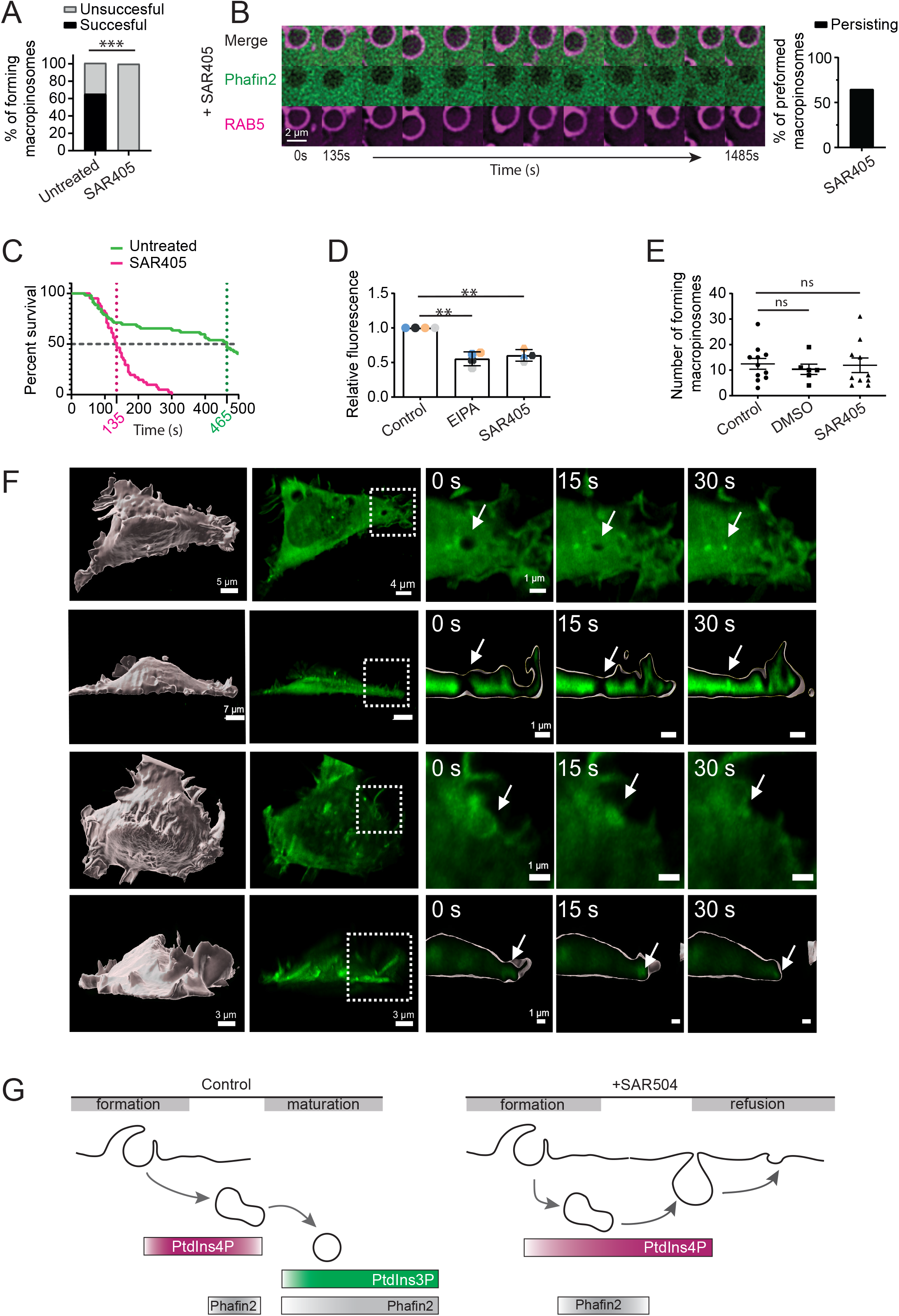
VPS34 inhibition induces re-fusion of newly formed macropinosomes with the plasma membrane (A) Success of macropinosome formation in the presence or absence of VPS34-inhibitor (3 μM) SAR405 was scored by live cell imaging. Newly forming macropinosomes were identified by the first, transient Phafin2 localization and then followed by either the Phafin2-GFP signal (control) or the dark, cytosol-excluding structures easily visible after SAR405 treatment. Whereas vesicles in untreated conditions were stable and could be observed for prolonged time periods, macropinosomes in SAR405-treated cells could be observed as dark, cytosol excluding structures that shrank and disappeared. In untreated conditions 63% of forming macropinosomes survived and reached the endosomal stage. Upon SAR405 treatment the number of macropinosomes surviving was reduced to 0%. (untreated events n= 53/ 9 cells; SAR 405 treated events n= 45/ 6 cells) Fisher’s exact test; two-sided, P<0.0001 (B) Vesicles with established endosomal identity are not affected by VPS34 inhibition. Shown is a macropinosome expressing Phafin2-GFP and mCherry-RAB5, treated with SAR405, which had already acquired RAB5 prior to addition of the inhibitor. The vesicle is stable over extended time periods and does not collapse. The majority (65%) of pre-formed, RAB5-labelled vesicles persisted throughout the duration of imaging after SAR405 treatment. (C) Lifetimes as an indication of macropinosome survival in control and SAR405-treated conditions are plotted (untreated events n= 52/ 9 cells; SAR 405 treated events n= 41/ 6 cells). Macropinosomes in both conditions were manually tracked and the track lengths were used as lifetimes. SAR405 treatment significantly reduces macropinosome lifetime compared to control. Log-rank test p<0.0001 (mean lifetime control = 465s; mean lifetime SAR405-treated: 135s). (D) Flow cytometry assay to determine 70 kDa dextran uptake in untreated and SAR405 treated cells. Cells that have been treated with SAR405 show reduced 70kDa dextran uptake in comparison to control cells. The macropinocytosis inhibitor EIPA was used as a positive control and shows significant reduction in dextran uptake (n=4 independent experiments, error bars: SD, statistic test: One-way ANOVA/ Dunnett’s multiple comparisons test: Control vs EIPA p=0.0051; Control vs SAR405 p=0.0042) (E) Effect of SAR405 on the number of initialized macropinosomes. Macropinosomes showing the first Phafin2 recruitment were counted under untreated, DMSO control and SAR405 treated conditions. There are no significant changes between the three condition in the number of forming macropinosomes (error bars: SEM, Control: n=9 cells/ 137 macropinosomes; DMSO: n=6 cells/62 macropinosomes; SAR405: n=9 cells/ 119 macropinosomes, One-Way ANOVA with Dunnet’s test: Control vs DMSO p= 0.8081; Control vs SAR405 p= 0.9801). (F) High resolution light sheet microscopy of HT1080 cells expressing Phafin2-mNeonGreen after SAR405 treatment. Shown are two example cells from a top view and a side view. The insets show macropinosomes which shrink and re-fuse with the plasma membrane. (G) Model of phosphoinositide turnover and dynamics on newly forming macropinosomes in the absence and presence of the VPS34-inhibitor SAR405.

We then asked what happened to the newly formed macropinosomes, which disappeared when VPS34 was inhibited. When performing 2D imaging, it appeared that macropinosomes shortly after formation started to rapidly shrink until they vanished (Figure 1E). The morphology and dynamics of these events suggested that they re-fused with the plasma membrane. To visualize this process, we performed high resolution oblique plane light sheet microscopy (OPM), which allows fast volumetric imaging with minimal photobleaching [32]. Using this method, we followed the fate of individual macropinosomes after inhibition of VPS34 with SAR405 in 3D. We observed that macropinosomes shrunk and often, a thin connection to the plasma membrane was visible, further indicating that VPS34 inhibition triggered re-fusion of nascent vesicles with the plasma membrane (Figure 2F, movie S2).

Our data indicates that failure to establish PtdIns3P leads to macropinosome refusion with the plasma membrane (Figure 2G). In contrast, already established macropinosomes that have gained endosomal identity – indicated by RAB5 - are stable and not affected by VPS34 inhibition.

### VPS34 inhibition blocks recruitment of endocytic factors

As we found that vesicles with an established endocytic identity are not affected by VPS34 inhibition, whereas newly forming vesicles are highly dependent on VPS34 activity, we reasoned that PtdIns3P might be required to recruit a specific factor, which controls endocytic identity. We therefore tested early endocytic markers and investigated if these are still recruited to the forming macropinosome.

One of the first factors recruited to forming macropinosomes is the RAB5 effector protein APPL1 [17]. In addition to RAB5 it binds to the membrane by a PH and a BAR domain but does not interact with PtdIns3P [33, 34]. In non-treated control cells, we observed recruitment of mCherry-APPL1 to macropinosomes; this localization occurs after the initial Phafin2 localization (Figure 3A) [23]. In contrast, VPS34-inihibition completely blocked recruitment of mCherry-APPL1 to forming macropinosomes (Figure 3A). This was surprising, as APPL1 recruitment is not dependent on PtdIns3P and APPL1 was not displaced from existing vesicles by inhibition of VPS34 (Supplementary Figure S1).

**Figure 3.**
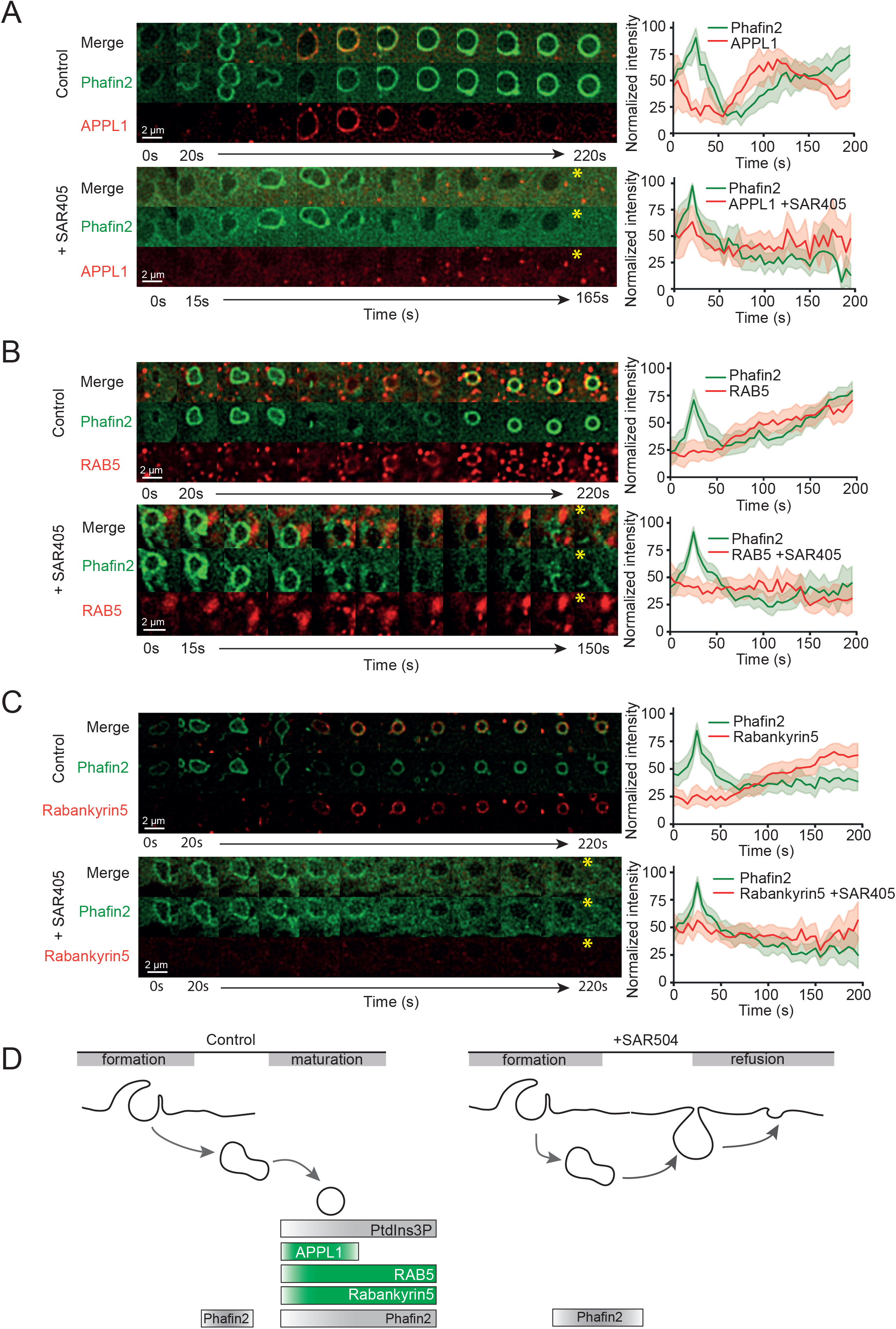
VPS34 inhibition blocks recruitment of endosomal factors to the newly formed macropinosomes. (A) Comparison of mCherry-APPL1 recruitment to forming macropinosomes in the absence and presence of SAR405. In control conditions, mCherry-APPL1 is transiently recruited to macropinosomes with the onset of the second Phafin2-GFP recruitment (n=22; mean + 95% CI). Upon VPS34-inhibiton mCherry-APPL1 recruitment to forming macropinosomes is abolished (n= 22; mean + 95% CI). Representative sequential images for both conditions are shown. Asterisk indicates end of macropinosome lifetime. (B) Comparison of mCherry-RAB5 recruitment to forming macropinosomes in the absence and presence of SAR405. In control conditions, mCherry-RAB5 is recruited to macropinosomes with the onset of the second Phafin2-GFP recruitment (n=45; mean +95% CI). Upon VPS34-inhibition mCherry-RAB5 recruitment to forming macropinosomes is abolished (n=38; mean + 95% CI). Representative sequential images for both conditions are shown. Asterisk indicates end of macropinosome lifetime. (C) Comparison of mCherry-Rabankyrin5 recruitment to forming macropinosomes in the absence and presence of SAR405. In control conditions mCherry-Rabankyrin5 is recruited to forming macropinosomes with the onset of the second Phafin2-GFP recruitment (n=31; mean+ 95% CI). In SAR405 treated conditions recruitment of Rabankyrin5 to forming macropinosomes is abolished (n=36; mean + 95% CI). Representative sequential images for both conditions are shown. Asterisk indicates end of macropinosome lifetime. (D) Schematic overview of the recruitment dynamics of the different endosomal proteins to forming macropinosomes in the absence and presence of SAR405.

Recruitment of APPL1 depends on RAB5, which is one of the key GTPases regulating early endocytosis and is involved in macropinosome formation and closure [35] as well as endosome fusion and maturation [36, 37]. RAB5 is further important for recruitment and regulation of the endosomal VPS34-complex [22, 37, 38]. We therefore tested if VPS34 inhibition would influence RAB5 recruitment. Using live cell imaging, we observed that mCherry-RAB5a was recruited to macropinosomes directly after the first transient Phafin2-GFP recruitment (Figure 3B). In contrast, VPS34 inhibition completely abolished the recruitment of mCherry-RAB5 to newly formed macropinosomes (Figure 3B). While new vesicles were still initiated and showed the first, transient Phafin2 localization, RAB5 was never recruited to the macropinosome, suggesting that PtdIns3P is critical to establish RAB5 on macropinosome membranes.

In line with this, the RAB5 effector and FYVE domain containing protein Rabankyrin5 also failed to be recruited. Rabankyrin5 localizes to macropinosomes and regulates endocytic trafficking [16, 23]. Under control conditions, mCherry-Rabankyrin5 localized to macropinosomes directly after the initial Phafin2 localization. In line with the absence of both RAB5 and PtdIns3P, VPS34 inhibition completely blocked this recruitment (Figure 3C).

Our findings that VPS34-inhibition completely abolishes recruitment of RAB5 and the RAB5-effectors APPL1 and Rabankyrin5 to newly forming macropinosomes, suggest that VPS34-generated PtdIns3P is essential for newly formed macropinosomes to recruit endosomal markers and establish endosomal identity (Figure 3D).

### VPS34 inhibition imparts a recycling identity on macropinosomes

Live-cell imaging (Figure 1E, 2F) suggested that nascent macropinosomes in SAR405-treated cells re-fused with the plasma membrane. As macropinosomes under VPS34-inhibited conditions failed to establish endocytic markers like RAB5, and harbored a PtdIns4P pool, we asked if these vesicles might have a secretory identity. Therefore, we examined if VPS34 inhibition affected recruitment of different RAB GTPases involved in recycling and secretion. To this end, we transfected different RAB GTPases involved in endocytic recycling using a Phafin2-GFP stable line and tracked macropinocytosis by live cell imaging.

First, we examined RAB8, which is involved in exocytic trafficking to the plasma membrane but has also been shown to localize to early macropinosomes [39–42].

When imaging Phafin2-GFP and mCherry-RAB8 under untreated conditions, we observed that mCherry-RAB8 localized to macropinosomes in the early stages of formation at the plasma membrane. mCherry-RAB8 recruitment coincided with the first Phafin2-GFP recruitment. mCherry-RAB8 localized transiently to macropinosomes and then dissociated with the onset of the second Phafin2-GFP recruitment (Figure 4A). Upon VPS34-inhibition, mCherry-RAB8 showed increased and persistent localization to macropinosomes. This localization lasted until the re-fusion of the macropinosome with the plasma membrane (Figure 4A). Next, we examined RAB11, which mediates slow recycling via the endosomal recycling compartment [43]. Under control conditions, we observed no recruitment of mCherry-RAB11 to forming macropinosomes (Figure 4B). However, after addition of VPS34-inhibitor, we observed a strong recruitment of RAB11 to forming macropinosomes (Figure 4B). The recruitment of mCherry-RAB11 upon VPS34 inhibition started with the peak of the Phafin2-GFP recruitment and persisted until the disappearance of macropinosomes. Next, we analyzed the behavior of RAB10, a small GTPase belonging to the same family as RAB8, which is also involved in endocytic recycling [44–46]. We observed that mCherry-RAB10 transiently accumulated on forming macropinosomes in untreated cells directly after the first Phafin2 recruitment (Figure 4C). In SAR405-treated cells, initial mCherry-RAB10 recruitment was unchanged, but mCherry-RAB10 showed a persistent accumulation on macropinosomes until they refused with the membrane (Figure 4C).

**Figure 4.**
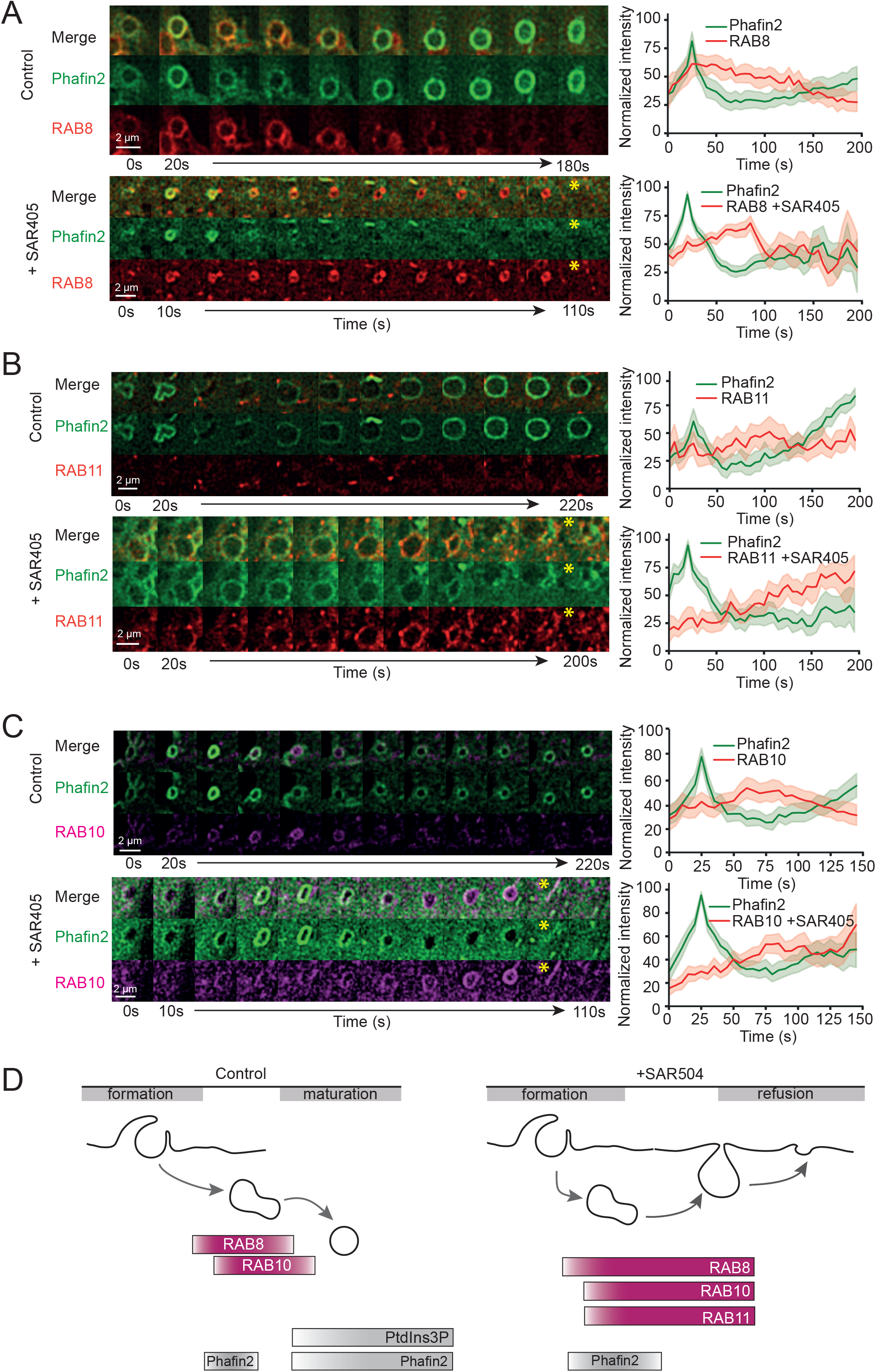
VPS34 inhibition leads to accumulation of secretory and recycling factors on newly formed macropinosomes. (A) Comparison of RAB8 recruitment to macropinosomes in the absence and presence of SAR405. In untreated conditions mCherry-RAB8 is recruited with the first Phafin2-GFP recruitment to newly forming macropinosomes and starts to disappear with the onset of the second Phafin2-GFP recruitment (n=33; mean + 95% CI). Upon VPS34-inhibition mCherry-RAB8 recruitment still occurs with the first recruitment of Phafin2-GFP, but then accumulates on macropinosomes until the end of their life (n=96; mean+ 95% CI). Representative sequential images for both conditions are shown. Asterisk indicates end of macropinosome lifetime. (B) Comparison of RAB11 recruitment to forming macropinosomes in the presence or absence of SAR405. In the absence of the VPS34-inhibitor mCherry-RAB11 is sparsely recruited to subdomains on forming macropinosomes (n=21; mean + 95% CI). Upon addition of SAR405 mCherry-RAB11 is recruited to macropinosomes with the decline of the first Phafin2-GFP recruitment and remains for the rest of the macropinosomes lifetime (n=25; mean + 95% CI). Representative sequential images for both conditions are shown. Asterisk indicates end of macropinosome lifetime. (C) Comparison of RAB10 recruitment to forming macropinosomes in the presence or absence of SAR405. In the absence of the VPS34-inhibitor mCherry-RAB10 is transiently recruited to forming macropinosomes (n=42; mean+ 95% CI). Upon addition of SAR405 mCherry-RAB10 is recruited to macropinosomes with the decline of the first Phafin2-GFP recruitment and remains for the rest of the macropinosomes lifetime (n=50; mean +95% CI). Representative sequential images for both conditions are shown. Asterisk indicates end of macropinosome lifetime. (D) Schematic overview of the recruitment dynamics of different recycling factors to forming macropinosomes in the absence and presence of SAR405.

Taken together, we find that VPS34 inhibition increases recruitment of RAB proteins involved in recycling and secretion, like RAB8, RAB11 and RAB10 on newly forming macropinosomes, suggesting that these vesicles gain a secretory identity (Figure 4D).

### RAB5 is critical for successful macropinosome formation

We next asked if the failure of forming macropinosomes to recruit RAB5 - and thus establish endosomal identity - is the cause for the re-fusion of the vesicle with the plasma membrane. To test this hypothesis, we depleted all three RAB5 isoforms (RAB5A, RAB5B, RAB5C) by siRNA and monitored macropinocytosis in control and knockdown cells. First, we tested if RAB5 depletion affects fluid phase uptake by measuring 10 kDa dextran internalization using flow cytometry. Both non-treated cells and siControl treated cells showed similar efficiency in dextran uptake. In contrast, either treatment with SAR405 or depletion of RAB5 (siRAB5ABC) led to a strong reduction in dextran uptake (Figure 5A,B).

**Figure 5.**
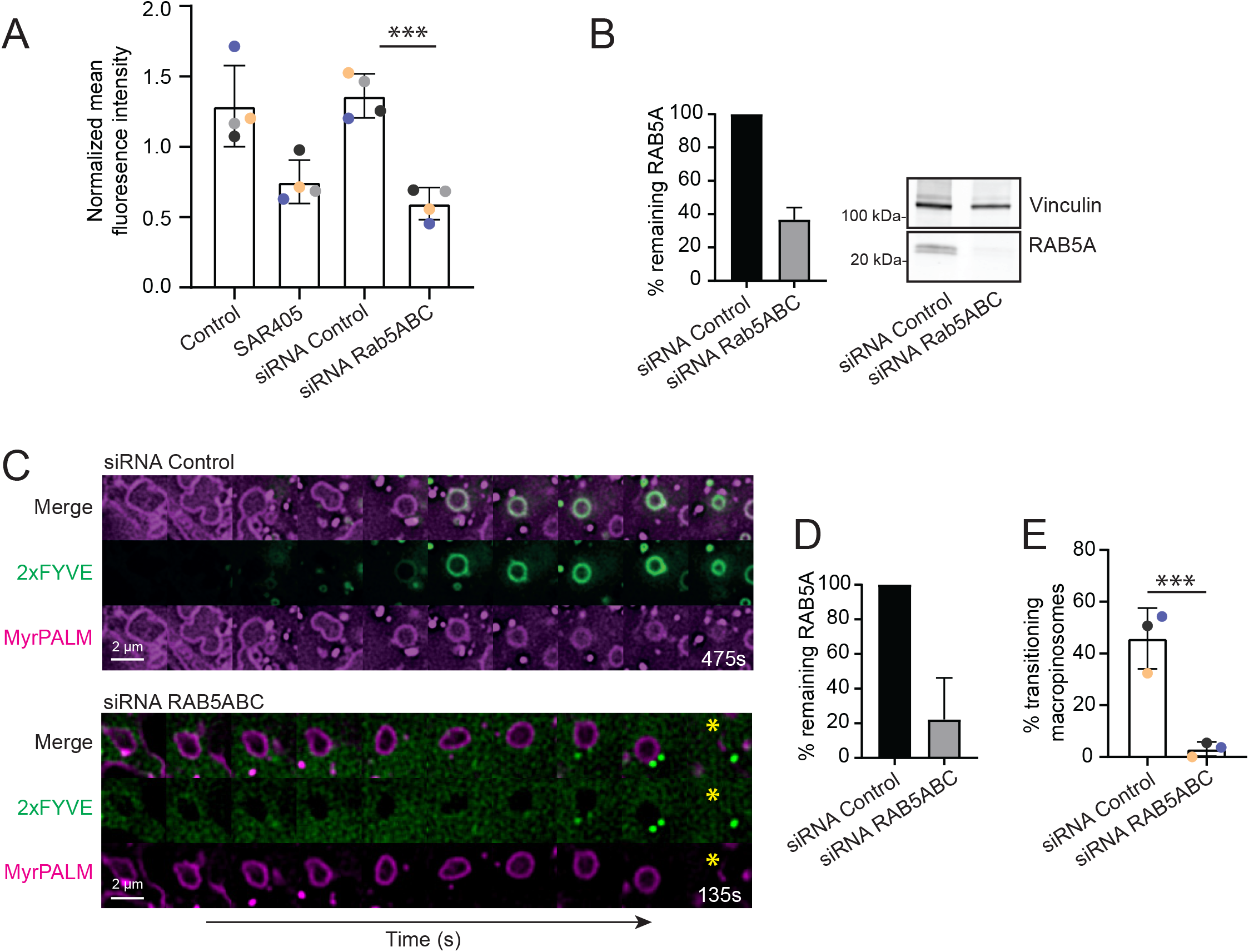
RAB5 recruitment to forming macropinosomes is critical for successful macropinosome formation. (A) Flow cytometry assay in RAB5 knock down cells measuring 10 kDa dextran internalization by macropinocytosis. The graph represents the relative fluorescence intensity per condition normalized to the mean intensity of all conditions. Error bars denote mean +/− SD from n=4 independent experiments indicated by different colors; * P<0.01, ***P<0.001, unpaired t-test siControl vs siRNA RAB5ABC p= 0.0002. (B) Knock down verification of RAB5 for the 4 independent dextran internalization experiments in Fig 5A. Representative western blots are shown. (C) Example galleries of MyrPALM positive macropinosomes that transition to the endosomal 2xFYVE stage or collapse at the MyrPALM positive stage. In the first gallery a successful transition from the early MyrPALM positive stage to the later, endosomal 2xFYVE positive stage is shown. In the second gallery a macropinosome that does not manage to acquire the 2xFYVE probe and instead collapses is shown. Asterisk indicates end of macropinosome lifetime. (D) Knock down verification of RAB5 for the 3 independent MyrPALM to 2xFYVE transition experiments. (E) Quantification of macropinosome transitions from MyrPALM to 2xFYVE in cells transfected with siRNA control or siRNA against RAB5ABC. The graph represents the % of macropinosomes that undergo transition. Error bars denote mean +/− SD from n=3 independent experiments indicated by different colors. siControl: 11 cells, 326 events; RAB5ABC KD: 14 cells, 248 events; siControl vs. siRNA RAB5ABC p= 0.0036, unpaired t-test, two tailed.

The role of RAB5 in macropinosome formation is not completely clear. While some studies place RAB5 already at the closure of the macropinosome and the scission from the plasma membrane [35], we see RAB5 only arriving once the macropinosome has been closed and after the initial Phafin2 recruitment [23]. We therefore asked if RAB5 knockdown cells could still initialize macropinosomes, or if already this step was perturbed. To track macropinosome formation, we used plasma-membrane localized mCherry (MyrPalm-mCherry) together with mNeonGreen-2xFYVE to mark the endosomal stage of macropinosomes. Using this assay, we tracked macropinosomes from the formation of cup-shaped membrane ruffles until they entered the endocytic pathway. Cells transfected with siControl showed normal formation of macropinosomes, with membrane ruffles forming cup-shaped, closed vesicles (outlined by MyrPalm-mCherry), which then gradually gained 2xFYVE, indicating that they accumulated PtdIns3P and established endosomal identity (Figure 5C,D). Likewise, cells depleted for RAB5 (siRAB5ABC) were able to form macropinosomes (Figure 5C,D). However, these vesicles did not gain 2xFYVE and ultimately re-fused with the plasma membrane. Tracking of individual macropinosomes showed that in control conditions, ~ 45 % of cup-shaped membrane ruffles transitioned to the endosomal stage, whereas less than 5% were able to do so in RAB5ABC knockdown cells (Figure 5E).

Thus, the lack of RAB5 shows the same phenotype as the lack of PtdIns3P, suggesting that the failure to recruit RAB5 in the absence of PtdIns3P could be the cause of the observed re-fusion of the macropinosomes with the plasma membrane.

### Forced recruitment of RAB5 does not rescue macropinosome survival upon VPS34 inhibition

Recent studies show that RAB5 not only recruits the VPS34 kinase complex, but that PtdIns3P enhances RAB5 membrane recruitment, which likely helps to activate RAB5 [47]. As we observed that RAB5 failed to be recruited to newly formed macropinosomes, we hypothesized that the initial recruitment and activation of RAB5 could be PtdIns3P dependent. In contrast to the dynamic wild-type RAB5, constitutively active RAB5 (RAB5Q79L) is permanently membrane associated once recruited [48, 49]. We therefore asked if expression of active RAB5 would be sufficient to overcome the effect of VPS34 inhibition. To this end, we expressed mCherry-tagged RAB5Q79L in cells stably expressing Phafin2-GFP (Figure 6A). Under control conditions, we observed robust recruitment of RAB5Q79L after the first Phafin2 recruitment, similar to wild-type RAB5 (Figure 6A). Addition of SAR405 did not affect RAB5Q79L recruitment, and treated cells showed a similar recruitment as non-treated cells (Figure 6A, movie S2). However, although these macropinosomes successfully recruited RAB5Q79L, this was not sufficient to prevent their re-fusion with the plasma membrane.

**Figure 6.**
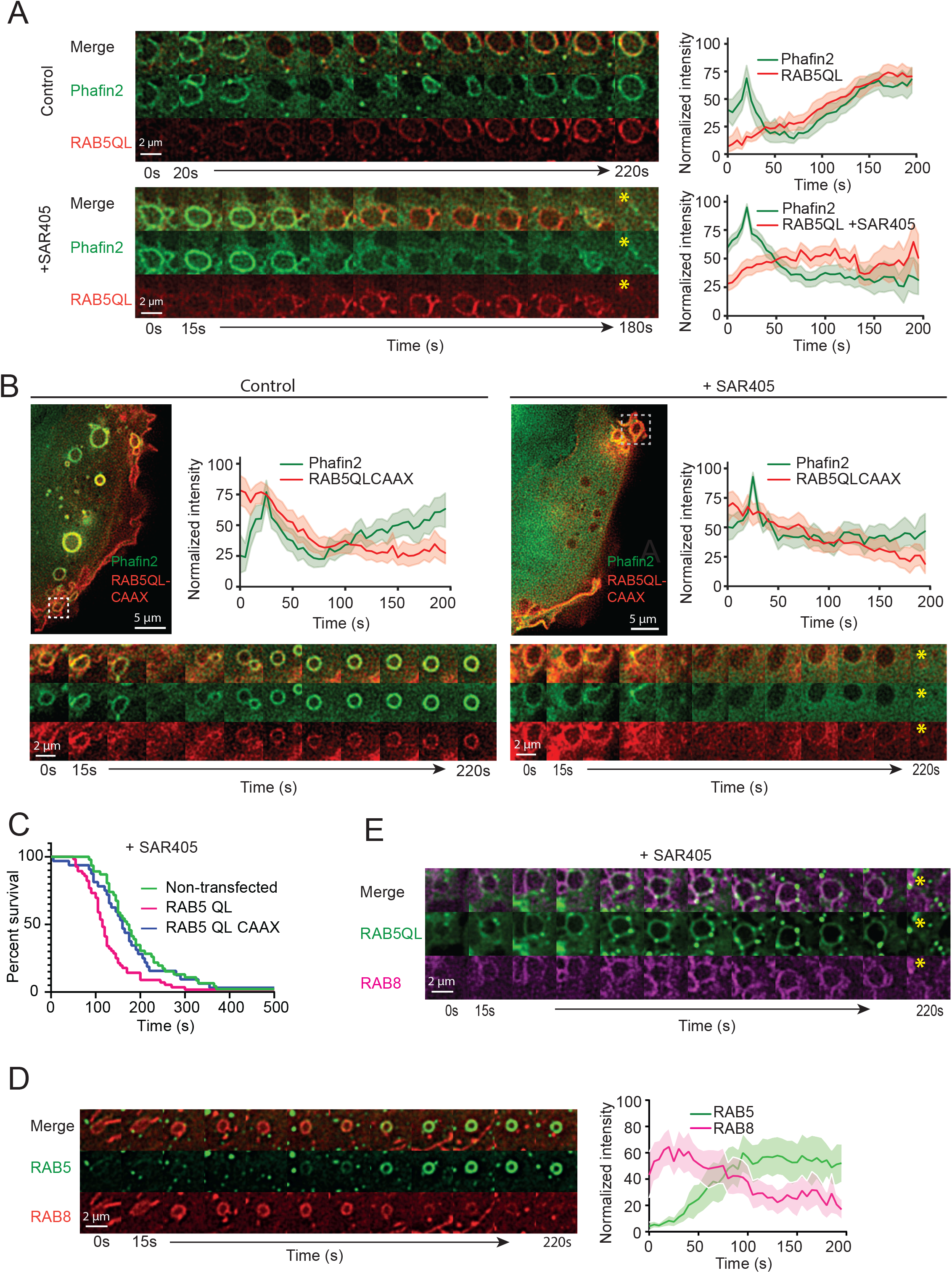
Forced recruitment of constitutively active RAB5 does not rescue macropinosome survival upon VPS34 inhibition. (A) Comparison of mCherry-RAB5 Q79L mutant recruitment in the absence or presence of SAR405. In control conditions RAB5 Q79L recruitment starts with the end of the first Phafin2-GFP recruitment and increases over time (n= 21; mean +95% CI). Upon VPS34-inhibition the constitutively active RAB5 Q79L mutant is still recruited to forming macropinosomes but does not restore macropinosome survival. Recruitment of mCherry-RAB5Q79L starts with the first Phafin2-GFP peak and continues until collapse of the macropinosomes (n=56; mean +95% CI). Representative sequential images for both conditions are shown (Phafin2-GFP, mCherry-RAB5 Q67L). Asterisk indicates end of macropinosome lifetime. (B) Comparison of mCherry-RAB5Q79L-KRas-CAAX recruitment in the absence or presence of SAR405. We exchanged the membrane targeting CXC domain of RAB5 against the plasma membrane targeting CAAX domain of KRas, to recruit RAB5 to macropinosomes at the plasma membrane. Performing live cell imaging we observe that mCherry-RAB5Q79L-KRas-CAAX is on forming macropinosomes with the onset of the first Phafin2-GFP recruitment and remains on macropinosomes during maturation (n=24; mean +95% CI). In SAR405 treated conditions mCherry-RAB5 Q79L-KRas-CAAX was still observed on forming macropinosomes with the onset of the Phafin2-GFP recruitment and remained there throughout the lifetime of the macropinosome (n= 32; mean +95% CI). Macropinosome survival was still reduced (asterisk). (C) Lifetimes of macropinosomes upon VPS34-inhibition in Phafin2-GFP cells with or without co-expression of RAB5Q79L or RAB5Q79L CAAX. Phafin2 + SAR405 events: 46; RAB5QL +SAR405 events: 56 macropinosomes; RAB5 QL CAAX +SAR405 events: 32 macropinosomes. (D) Macropinosomes were manually tracked in cells expressing both RAB8A and RAB5A. In the absence of Phafin2 as a macropinosome marker, we used RAB8 to track early forming macropinosomes. RAB8 localizes to macropinosomes at the ruffle stage. Once macropinosomes have been internalized they can be tracked by their exclusion of cytosol, as big dark spots. RAB8 localizes to macropinosomes in very early stages and then gradually decreases. The decrease of RAB8 is accompanied by an increase in RAB5 recruitment. The normalized intensity of each channel was plotted (n=16, error bars: mean + 95% CI). Representative sequential images of the recruitment dynamics of RAB8 and RAB5 to newly forming macropinosomes are shown. (E) Representative sequential images showing chimeric macropinosomes containing both RAB5Q79L and RAB8.

Several studies have placed RAB5 early in macropinocytosis, in some studies as early as during ruffle closure [16, 35, 50]. As we observed that RAB5Q79L recruitment occurred only after the initial Phafin2 recruitment, we wanted to exclude that this recruitment was too late to rescue macropinosome survival in SAR405 treated cells. We therefore exchanged the C-terminal CXC motif of RAB5 with the polybasic region and CAAX box of KRAS. This motif mediates robust plasma membrane recruitment [51]. Live cell imaging showed strong plasma membrane localization of RAB5Q79L-CAAX in both the presence and absence of SAR405 (Figure 6B). However, even the expression of these constructs was unable to rescue the macropinosome survival phenotype in SAR405-treated cells (Figure 6B). Quantification of macropinosome survival showed that expression of RAB5Q79L or RAB5Q79L-CAAX did not affect the average survival time of newly-forming vesicles upon SAR405 treatment (Figure 6C). This was surprising, as we found that vesicles with established endosomal identity were only mildly affected by SAR405 (Figure 2B).

### The transition from RAB8 to RAB5 requires VPS34 activity

As we found that SAR405 treatment led to the recruitment of secretory GTPases such as RAB8, RAB10 and RAB11 on newly-formed macropinosomes, we asked if SAR405 would establish a secretory identity on these vesicles even in the presence of RAB5. In non-treated cells expressing wild-type RAB8 and wild-type RAB5, we found a RAB switch, where RAB8 was dissociating, whereas RAB5 was associating with the newly-formed macropinosome (Figure 6D, movie S3). Intriguingly, in SAR405 treated cells, we observed robust and persistent localization of RAB8, even when the macropinosome in parallel recruited RAB5Q79L (Figure 6E, movie S4), and the vesicle re-fused with the plasma membrane. This suggests that the presence of active RAB5 is not sufficient to fully impart endocytic identity to macropinosomes in the absence of PtdIns3P, as these vesicles still acquire secretory characteristics. This implies that PtdIns3P could function to either recruit a yet unknown determinant of endosomal identity, or a displacement factor for RAB8 and other secretory GTPases, to stabilize the endocytic character of the newly-formed vesicle.

### Loss of PtdIns3P induces RAB8 and RAB10 mediated recycling of macropinosomes

In order to test if RAB8 drives the re-fusion of newly-formed macropinosomes with the plasma membrane, we depleted both isoforms of RAB8 by siRNA and analyzed the formation of macropinosomes. Indeed, in contrast to cells treated with control siRNA, we observed an increased success rate of macropinosome formation in cells depleted for RAB8A and RAB8B (Figure 7AB). On the other hand, overexpression of RAB8 reduced the number of successfully formed macropinosomes (Figure 7C). We also knocked down the related GTPase RAB10, as well as both RAB8 and RAB10, and assayed fluid phase uptake by flow cytometry (Figure 7D-F). In both cases, we observed enhanced fluid phase uptake in comparison to control siRNA (Figure 7D). These results are consistent with a role of RAB8 and RAB10 in recycling of macropinosomes.

**Figure 7.**
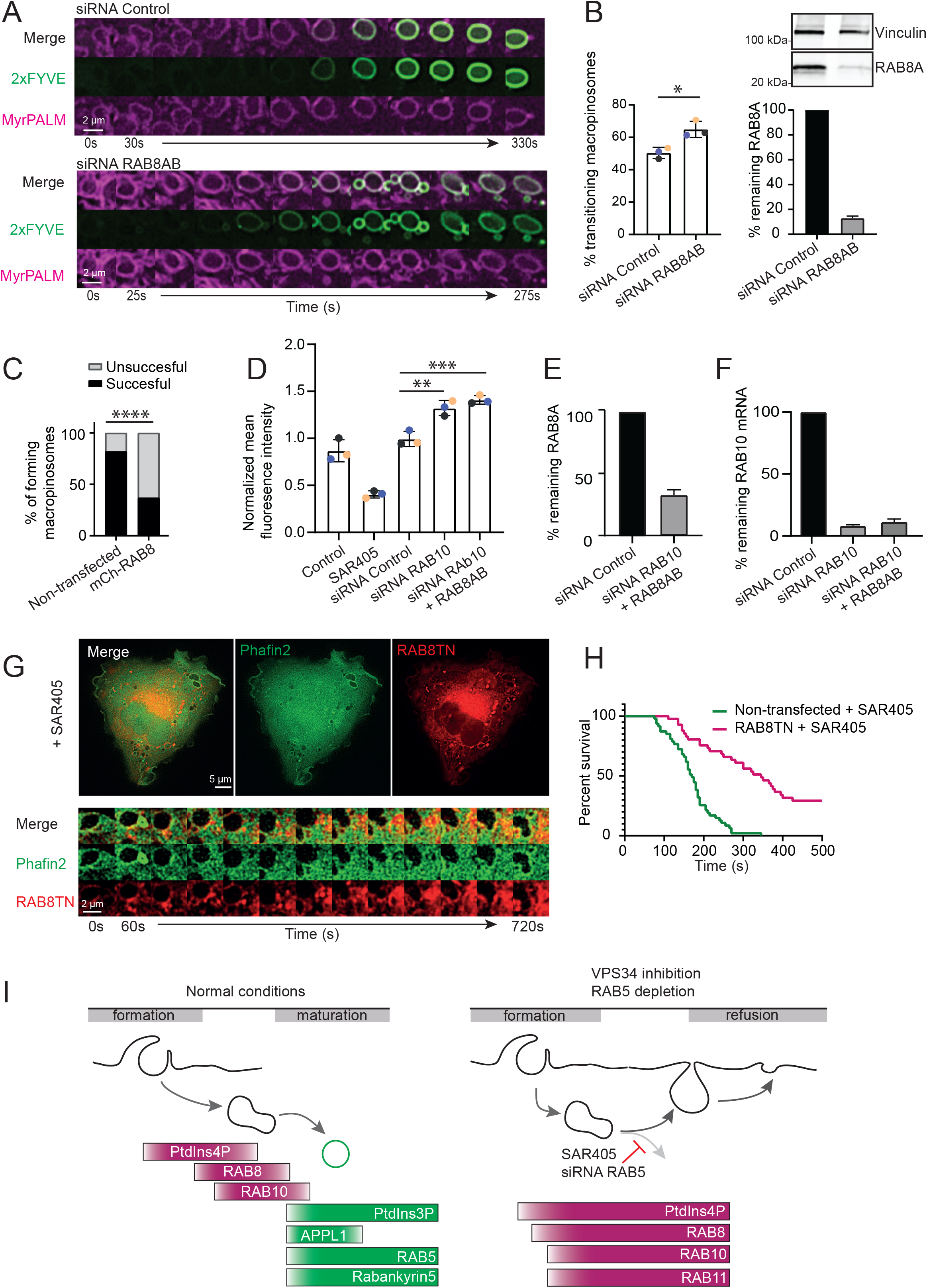
Newly formed macropinosomes undergo a VPS34-dependent transition from RAB8 to RAB5. Inhibition of this transition leads to RAB8 and RAB10 mediated plasma membrane re-fusion of newly formed macropinosomes. (A) Example galleries of MyrPALM positive macropinosomes that transition to the endosomal 2xFYVE stage or in control RNA or RAB8AB siRNA treated cells. In the first gallery, a successful transition from the early MyrPALM positive stage to the later, endosomal 2xFYVE positive stage is shown. In the second gallery, a successful transition from the early MyrPALM-positive stage to the 2xFYVE stage in RAB8AB depleted cells is shown. (B) Quantification of macropinosome transitions from MyrPALM to 2xFYVE in cells transfected with siRNA control or siRNA against RAB8AB. The graph represents the % of macropinosomes that undergo transition. Error bars denote mean +/− SD from n=3 independent experiments indicated by different colors. siControl: 15 cells, 353 events; RAB8AB KD: 13 cells, 349 events; siControl vs RAB8AB p= 0.0145; unpaired t-test, two tailed. Knock down verification of RAB8AB for the 3 independent MyrPALM to 2xFYVE transition experiments with representative western blots are shown. (C) Effects of overexpression of Phafin2 and RAB8WT on the success rate of forming macropinosomes. Macropinosomes were scored as successfully formed when the second Phafin2-GFP recruitment could be observed. In control conditions with only the overexpression of Phafin2 82% of macropinosomes formed successfully. Overexpression of RAB8WT decreased the success rate of newly forming macropinosomes to 37% (Chi^2^ test p< 0.0001). (D) Flow cytometry assay in RAB10 and RAB8 knock down cells measuring 10 kDa dextran internalization by macropinocytosis. The graph represents the relative fluorescence intensity per condition normalized to the mean intensity of all conditions. Error bars denote mean +/− SD from n=3 independent experiments indicated by different colors; ** P<0.01, ***P<0.001, one-way Anova with Dunnett’s correction, siControl vs RAB10 p= 0.0022; siControl vs RAB10+RAB8 p= 0.0007. (E) Knock down verification of RAB8 for the 3 independent dextran internalization experiments in Fig 7C. (F) Knock down verification of RAB10 using rt-qPCR for the 3 independent dextran internalization experiments in Fig7C. (G-H) Lifetime defects of macropinosomes caused by VPS34-inhibition are restored upon overexpression of RAB8 T22N. Representative image and gallery of RAB8 T22N localization to macropinosomes upon SAR405 treatment. Macropinosomes in the different conditions were manually tracked and the track lengths were used as lifetimes. Treatment with SAR405 significantly reduced lifetimes of newly forming macropinosomes. Overexpression of RAB8 T22N restored the lifetimes of newly forming macropinosomes in the presence of SAR405 (Kruskal-Wallis, Dunn’s test SAR405 vs RAB8 T22N SAR405 p< 0.001). Phafin2 + SAR405 events: 47; Phafin2 + RAB8 T22N events: 41. (I) Under normal conditions, newly formed macropinosomes undergo a transition from a secretory identity to an endosomal identity. Initially newly formed macropinosomes are positive for PtdIns4P and secretory RABGTPases like RAB8 and RAB10. With the onset of VPS34-activity and PtdIns4P dephosphorylation to PtdIns by phosphatases, PtdIns3P begins to accumulate on the macropinosome membrane. This destabilizes the secretory RABGTPases and their effectors and allows recruitment of endosomal factors like RAB5 and its effectors. Since active RAB5 enhances VPS34-recruitment and PtdIns3P increases RAB5 activation, a positive feedback loop is generated, which stably establishes macropinosomes in the endo-lysosomal pathway. When we inhibit VPS34 or deplete RAB5, newly formed macropinosomes cannot undergo their phosphoinositide transition. Instead of accumulating PtdIns3P, PtdIns4P is stabilized on the macropinosome membrane, allowing accumulation of factors like RAB8, RAB10 and RAB11. This accumulation of secretory factors stably establishes the newly formed macropinosomes in the recycling pathway resulting in their re-fusion of the plasma membrane.

We then proceeded to test if inhibition of RAB8 activity could suppress the effect of VPS34 inhibition and prevent re-fusion of nascent macropinosomes with the plasma membrane. As depletion of RAB8 by siRNA was not complete, we chose to express dominant negative RAB8T22N instead. This mutant is unable to bind GTP, which often results in more robust phenotypes than siRNA-mediated depletion. We transfected cells expressing Phafin2-GFP with mCherry-RAB8T22N, treated them with SAR405 and analyzed macropinosome formation and survival by live cell microscopy (Figure 7G,H). In control cells, SAR405 treatment led to rapid re-fusion of nascent macropinosomes with the plasma membrane. In contrast, expression of RAB8T22N stabilized newly-forming macropinosomes even in the absence of PtdIns3P and suppressed the re-fusion with the plasma membrane (Figure 7G,H, movie S5).

Taken together, our results show that VPS34 regulates the balance between macropinsome maturation and exocytosis by controlling the dynamic transition of PtdIns4P to PtdIns3P, and RAB8 to RAB5 (Figure 7I).

## Discussion

In this study, we provide insight into the mechanisms that establish the membrane identity of newly-formed vesicles by using macropinosomes as a model system. Under normal conditions, macropinosome maturation is characterized by a change in PIs from PtdIns4P to PtdIns3P in concert with a transition of RAB8 to RAB5. When VPS34 is inhibited, macropinosomes accumulate RAB8 and other recycling factors, whereas RAB5 recruitment and endocytic maturation is prevented. This leads to re-fusion of the macropinosome with the plasma membrane (Figure 7I). Since the forced recruitment of RAB5 to SAR405 treated macropinosomes failed to rescue macropinosome survival, this indicates that VPS34 has a dual role in macropinosome maturation. In addition to enabling the recruitment of RAB5, VPS34 is also required for the removal of RAB8. Such a mechanism could ensure that forming macropinosomes will mature through the endosomal pathway to deliver their cargo for degradation in lysosomes, rather than being recycled and exocytosed.

How can VPS34 control the fate of newly-formed vesicles? Early endocytic vesicles carry RAB5, which recruits effector proteins that define the biochemical properties of the vesicle. In addition, these vesicles gradually gain PtdIns3P, which can then recruit other effector proteins and ultimately controls the maturation of the endocytic vesicles to early and late endosomes [28, 37]. RAB5 binds to the VPS34 kinase complex, which generates PtdIns3P and regulates its activity [38, 52]. At the same time, PtdIns3P enhances membrane recruitment of RAB5 [47] and could potentially form a positive feedback loop. PtdIns3P also drives the maturation of early to late endosomes and the switch of RAB5 to RAB7 [53–55].

We find that PtdIns3P – generated by the VPS34 kinase complex – is not only required for the maturation of early to late endosomes but is also critical for the initial establishment of endosomal identity. Inhibition of VPS34 – and consequently the absence of PtdIns3P – completely blocks the recruitment of RAB5 to newly formed macropinosomes and prevents these vesicles from gaining endosomal identity.

RAB5 recruitment and activation is a fine-tuned system that heavily relies on the guanine nucleotide exchange factor (GEF) Rabex5, complexed to the RAB5 effector Rabaptin5 as well as the endosomal VPS34-complex. PtdIns3P could act as an initial recruiter of RAB5 [47, 56]. Membrane-associated RAB5 could be activated by Rabex5, and active RAB5 would then recruit more Rabex5-Rabaptin complex, thus driving a positive feedback loop [57]. Active RAB5 would also recruit the endosomal VPS34 complex and thereby produce more PtdIns3P, further promoting the recruitment of RAB5. The gradual increase of PtdIns3P at endosomal membranes supports such a positive feedback model.

The inhibited recruitment of RAB5 and the RAB5 effectors APPL1 and Rabankyrin5 that we observed in response to VPS34 inhibition could be due to the inability of RAB5 and its activation complex to overcome this initial hurdle to establish active RAB5 domains without active VPS34. This would also fit with our observation that constitutively active RAB5Q79L can rescue RAB5 localization to newly formed macropinosomes under VPS34-inhibited conditions.

This model would imply that either an initial pool of PtdIns3P or a stochastic activation of RAB5 is needed to overcome the initial inertia and establish active RAB5 domains on macropinosomes. Indeed, previous studies report a transient, phosphatase derived PtdIns3P pool which could fulfill this role. A phosphatase cascade on macropinosomes can metabolize PtdIns3,4,5P3 via PtdIns3,4P2 to PtdIns3P [14]. It is tempting to speculate that this PtdIns3P pool could act as an initial trigger for RAB5 recruitment. Alternatively, stochastic membrane association and activation of RAB5 might be sufficient to activate VPS34 and thereby initiate recruitment of RAB5.

However, active RAB5 is not sufficient to overcome the need for PtdIns3P, since neither constitutive active RAB5Q79L, nor membrane-tethered RAB5Q79L-CAAX, which both are localizing to newly formed macropinosomes, prevent the re-fusion of macropinosomes in the absence of active VPS34. We find that PtdIns3P, produced by VPS34, has a dual function. Apart from the recruitment of RAB5, it is also needed to prevent nascent macropinosomes from acquiring a secretory identity.

We show that macropinosomes undergo a shedding of their plasma membrane identity, by undergoing a RAB and phosphoinositide switch from a RAB8 and PtdIns4P positive stage to a RAB5 and PtdIns3P positive stage. This corresponds to a switch from a recycling identity – as PtdIns4P and RAB8 are key regulators of secretion – to an endocytic identity. VPS34 activity appears to be a critical regulator of this switch, as in the absence of PtdIns3P, RAB8 and PtdIns4P remain on the macropinosome and ultimately drive its refusion with the plasma membrane. Similarly, other recycling factors – such as RAB11 and RAB10 – are recruited if VPS34 is inhibited. Recycling and re-fusion with the plasma membrane can be blocked by overexpression of dominant negative RAB8 T22N, suggesting that the prolonged recruitment of RAB8 is one of the major drivers of macropinosome re-fusion. Our findings also suggest that PtdIns3P is required to remove RAB8 and RAB10 from the macropinosome and thus remove this secretory identity.

Interestingly, a recent study also described a re-fusion of macropinosomes with the plasma membrane [35]. This study shows that RAB5-mediated recruitment of OCRL and INPP5b via APPL1 is required to remove PtdIns4,5P2 from macropinosomes. Removal of PtdIns4,5P2 is critical to prevent re-fusion with the plasma membrane [35]. If VPS34 is inhibited, the failure to establish RAB5 at the limiting membrane of macropinosomes could similarly result in a failure to recruit OCRL and INPP5b, which could cause the re-fusion observed by us and others [35]. Both OCRL and INPP5b are 5’ phosphatases which could contribute to the production of the transient PtdIns4P pool we observe. While this is an attractive model, it does not completely match our findings. First, both OCRL and INPP5b act at very early stages of macropinosome formation, during or directly after the scission from the membrane, whereas we observe that vesicles are formed, but fail to establish endosomal identity. Moreover, we find that the transient PtdIns4P pool is not affected if we inhibit VPS34, making it unlikely that RAB5-controlled OCRL and INPP5b activity is responsible to generate this pool. Moreover, RAB5 Q79L – which should recruit APPL1 and by proxy also OCRL and INPP5b to forming macropinosomes, does not suppress the collapse of macropinosomes.

An attractive alternative model is that VPS34 actively competes with PI4 kinases for PtdIns. In the absence of active VPS34, PI4 kinases could metabolize PtdIns on the newly formed macropinosome, priming them for recycling. A similar mechanism has been described for recycling endosomes. On endosomes, the phosphatase MTM1 controls hydrolysis of PtdIns3P to PtdIns [58]. PtdIns then serves as substrate for PI4K2α, which generates PtdIns4P [58]. PtdIns4P is a cue for secretory identity and recruits the exocyst complex, whereas accumulation of PtdIns3P and its effectors prevent recycling [58]. These findings have several similarities to the maturation of macropinosomes. As mentioned above, a critical step during macropinosome internalization is the dephosphorylation of PtdIns3,4,5P3 to PtdIns. One of the involved phosphatases - MTMR6 – dephosphorylates phosphatase-derived PtdIns3P to PtdIns, which could then be phosphorylated to PtdIns4P [14]. VPS34 could counter-balance this process and limit the accumulation of PtdIns4P by generating PtdIns3P. Together with the RAB5-driven positive feedback loop, this would generate a robust endocytic identity. Thus, generation of PtdIns3P on macropinosomes by VPS34 could limit available PtdIns4P and the stabilize the endocytic identity.

Taken together, we show in this study that forming macropinosomes need to undergo a transition from RAB8 to RAB5 in order to mature along the endo-lysosomal pathway, similar to the transition from early endosomes marked by RAB5 to late endosomes and lysosomes marked by RAB7 [53]. This transition is functionally required for successful macropinosome maturation and is regulated by phosphoinositide transitions and gradients that allow a tightly controlled recruitment of effector proteins needed for the different stages. The default secretory identity of macropinosomes could be a fail-safe mechanism that prevents the spurious uptake of vesicles and requires that they actively gain endocytic activity. This would prevent the accumulation of “vagrant” vesicles lacking a defined identity. At the same time, it could provide a first layer of defense against pathogen invasion, as a pathogen entering cells by induced macropinocytosis would have to actively modulate the vesicle coat or end up in either the endolysosomal pathway or the secretory pathway [59].

We propose the following model for endocytic macropinosome maturation: Newly forming macropinosomes have a secretory identity by default. In order to mature along the endo-lysosomal pathway, they need to be stripped of secretion promoting lipids such as PtdIns4P and RABGTPases, such as RAB8 and RAB10. Maturation along the endo-lysosomal pathway requires both PtdIns3P, RAB5 and RAB5 effectors. Together VPS34 and RAB5 create a stabilizing feedback loop, which allows for the generation of more PtdIns3P and in turn displacement of RAB8 and RAB10 as well as recruitment of RAB5 and RAB5 effectors, stably establishing macropinosomes in the endo-lysosomal pathway. Depleting either PtdIns3P or RAB5 inhibits entry into the endo-lysosomal pathway and instead stabilizes recycling identity on the newly formed macropinosomes, by recruitment of RAB8, RAB11 and RAB10 and PtdIns4P. Since entry into the endosomal pathway is blocked, these factors mediate re-fusion of the newly formed macropinosomes with the plasma membrane in order to maintain membrane homeostasis.

While our understanding of the exact regulation of the RAB8 to RAB5 transition is not yet complete, our findings elucidate how macropinosome maturation is regulated. This knowledge can potentially be utilized for investigating new treatment approaches for pathogen infections or cancers driven by macropinocytosis.

## Methods

### Cell lines

Experiments were performed in hTert-RPE1 cells (ATCC^®^ CRL-4000™) and HT1080 cells (ATCC^®^ CCL-121™). Cells were purchased from ATCC and authenticated by genotyping. Cells were verified to be free of mycoplasma contamination and regularly tested after manipulation.

Stable hTert-RPE1 or HT1080 cell lines were lentivirus generated pools using a third-generation system [60]. Tagged versions of proteins of interest were subcloned into a Gateway entry vector with a PGK or CMV promotor by standard molecular biology techniques. Transfer vectors were generated by a Gateway LR recombination with a destination vector of choice. Virus was packaged using a third-generation system, by co-transfecting a packaging vectors encoding for GAG/POL, REV and a VSV-G based envelope in addition to the destination vector. Viral supernatant was collected 72h after transfection. Cells were transduced with low viral titers (MOI<1) and stable pools were generated through antibiotic selection.

The hTert-RPE1 line stably expressing GFP tagged Phafin2 was generated previously [23] and used as a background for further stable lines. In this study, the following stable cell lines were used: hTert-RPE mNeonGreen-Rab5, hTert-RPE1 mCherry-Rab8, hTert-RPE1 mNeongreen-2xFYVE-MyrPalm-mCherry and HT1080 Phafin2-mNeonGreen-MyPALM-mCherry.

### Cell Culture

hTert-RPE1 cells were maintained in DMEM-F12 medium (Gibco) supplemented with 10% fetal bovine serum (Merck Life Science (Sigma)) and 5 U ml^−1^ penicillin and 50 μg ml^−1^ streptomycin (Merck Life Science (Sigma)) at 37 °C and 5% CO_2_. HT1080 cells were maintained in DMEM (Sigma Aldrich) supplemented with 10% fetal bovine serum and 5 U ml^−1^ penicillin and 50 μg ml^−1^ streptomycin at 37 °C and 5% CO_2_.

### Antibodies

The following primary antibodies were used: anti-RAB5A, Santa Cruz Biotechnology (sc-46692). Recombinant anti-RAB8A, Abcam (ab188574). Monoclonal anti-Vinculin, Sigma Aldrich/ Merck (V9131). Monoclonal anti-γ-Tubulin Sigma Aldrich/ Merck (T6557). Secondary antibodies were from LI-COR.

### Plasmids

The following plasmids were obtained from Addgene: pEGFP-RAB10 (Addgene #49472) was a gift from Marci Scidmore [61] and recloned into a pmCherry-C1 backbone. GFP-2xSidM (Addgene plasmid #51472) was a gift from Tamas Balla, GFP was exchanged with mCherry. pEGFP-APPL1 (Addgene plasmid #22198) was a gift from Pietro de Camilli [62], EGFP was exchanged with mCherry. pmCherry-RAB11A was a gift from Ellen M. Haugsten/ Jim Norman.

All other plasmids were generated using standard molecular cloning techniques.

### siRNA knockdown

siRNAs against RAB5A(S100301588), RAB5B (S102662800), RAB5C(S102663073), RAB8A (SI02662254), RAB8B(SI02662261), RAB10 (SI00301546, SI03037076) and non-targeting control were purchased from Qiagen AS (FlexiTube siRNA). Cells were transfected at around 50% confluency with Lipofectamine RNAiMax transfection reagent (Thermo Fisher) according to manufacturer’s instructions. Transfected cells were analyzed 48h after transfection.

### Flow Cytometry

Fluid phase uptake was measured using flow cytometry.

Briefly, cells were stimulated with 0,05μg/ mL HGF (Merck Life Science(Sigma) H5791) and incubated with 70 kDa Texas Red dextran (Molecular Probes/ Life Technologies D-1864) or 10kDa Alexa Flour 647 dextran (Thermo Fisher Scientific; D22914) for 15 min at 37 °C in the presence or absence of 6 μM SAR405 (Selleckchem S7682) in DMSO. When treated with EIPA (Merck Life science (Sigma) A3085), cells where pre-incubated with 50 μM EIPA for 30 min before adding dextran. Cells were washed 4-5 times with big amounts of pre-warmed medium and detached with Trypsin for 2-3min. Detached cells were measured on an LSR II flow cytometer using the 561 nm laser line.

### Transient Transfections

hTert-RPE1 cells were transfected using FuGENE 6 (Promega) according to the manufacturer’s instructions at a ratio of 1:3. Cells were grown in a 35 mm glass bottom dishes (MatTek Corporation) and transfected with up to 2 μg of DNA 12-14h before imaging.

### Live Cell Imaging

Cells were imaged in Life Cell Imaging buffer (Invitrogen) supplemented with 20 nM Glucose (Merck 1.08342.1000) and stimulated with 0,05 μg/ mL HGF (Merck Life Science (Sigma)) with or without 3 μM SAR405 (Selleckchem) in DMSO.

Live Cell imaging was performed on a Deltavision OMX V4 microscope (GE Healthcare) equipped with an Olympus 60x NA 1.42 objective, three PCO.edge sCMOS cameras, a solid-state light source and a laser-based autofocus. Environmental control was provided by a heated stage and an objective heater (20-20 Technologies). Cells were imaged in two or three colors in conventional mode every 5 seconds between 12-25 minutes in total. Acquired images were aligned and deconvolved using softWoRx software (Applied Precision, GE Healthcare) and x-y alignment was checked and when necessary re-calibrated using the “GE Image Registration slide”.

### OPM light sheet microscopy

Light sheet microscopy was performed using a custom-built oblique plane microscope. The microscope layout followed the general plan for a stage scanning OPM microscope described in Sapoznik et al [32]. The microscope was built around the ASI modular microscope platform. The inverted microscope consisted of a ASI MIM microscope body, an ASI FTP Z stage and a stage scanning optimized ASI X/Y stage (ASI imaging). Environmental control was provided by an Okolabs stage incubator. A 60x 1.3 NA silicon oil immersion lens (Olympus) served as primary objective. ASI cage elements were used to construct the rest of the optical train, consisting of a 300mm tube lens (TL1), a 357mm tube lens (Tl2) and a 40x 0.95 NA air remote objective (Nikon). The image generated by the remote objective was collected by a bespoke tertiary objective (AMS-AGY v1.0, Special Optics) and focused by a 250 mm tube lens (TL3) on the sensor of a Andor Zyla 4.2 camera, resulting in a final pixelsize of 91 nm. The excitation beam was coupled into the light path by a dichroic mirror placed between TL2 and the remote objective. We used a gaussian light sheet generated by an ASI light sheet generator (ASI Imaging) equipped with a cylindrical lens, coupled to a Toptica laser source (Toptica) with 405, 488, 561 and 647 nm laser lines. Image acquisition was performed by stage scanning. All hardware was synchronized by an ASI Tiger controller (ASI Imaging) and controlled by the ASI diSPIM plugin in MicroManager [63]. Deconvolution and deskewing were performed using the LLSPy software package (https://github.com/tlambert03/LLSpy/), the resulting images were visualized using Imaris.

### Quantitative real-time PCR of mRNA expression

mRNA expression analysis after siRNA transfection was performed as previously described in Pedersen et al. 2020 [64]. The primers used in this paper were obtained from Qiagen and are QT00067767 for RAB10 and QT00000721 for TATA-binding protein (TBP) as a reference housekeeping gene.

### Image processing and data analysis

All live cell imaging files were deconvolved using SoftWorx Software (GE Healthcare) before being analyzed and prepared for presentation. Analysis of life cell imaging was performed using Fiji and custom-made Python scripts (https://github.com/koschink/Phafin2). The intensity tracks were plotted using the Python Seaborn library. Newly forming macropinosomes were tracked manually in Fiji using Phafin2 as a marker for macropinosomes. When tracking macropinosomes in the absence of Phafin2-GFP, we used other transfected fluorescently tagged proteins, as well as the exclusion of cytosol from the big vesicular structures to follow the progress of the macropinosomes. The limiting membrane of the tracked vesicle was marked in each frame and added to the ROI manager. From the ROI’s the program measured the fluorescence intensity in every channel and created galleries for the tracked macropinosomes. Processing of the fluorescence intensity was performed by a Python script, which aligned the tracks to the first Phafin2-GFP peak.

### Statistics

No statistical methods were used to pre-determine sample size. All statistical calculations were derived from at least 3 biological replicates. For microscopy-based assays, each biological repeat included sufficient cell numbers to ensure that the effects could be robustly measured and was repeated at least 3 times. The number of experimental days, cells and vesicles analyzed are indicated in the figure legends. Data was tested for normal distribution using GraphPad Prism Version 8. For comparison of two samples with each other unpaired *t*-test was used for normally distributed data, while the Mann-Whitney test was used for non-normal data sets. For multiple comparisons we used the one-way analysis of variance (ANOVA) or the Kruskal-Wallis test with a suitable post hoc test. One-sample t-test was used when the value of control sample was set to 1. All error bars give the mean values ± 95% CI or as indicated in the figure legends. Samples were not randomized for this study.

### Data Availability

Source data for the figures is provided with the paper.

---

**Supplementary Figure 1.**
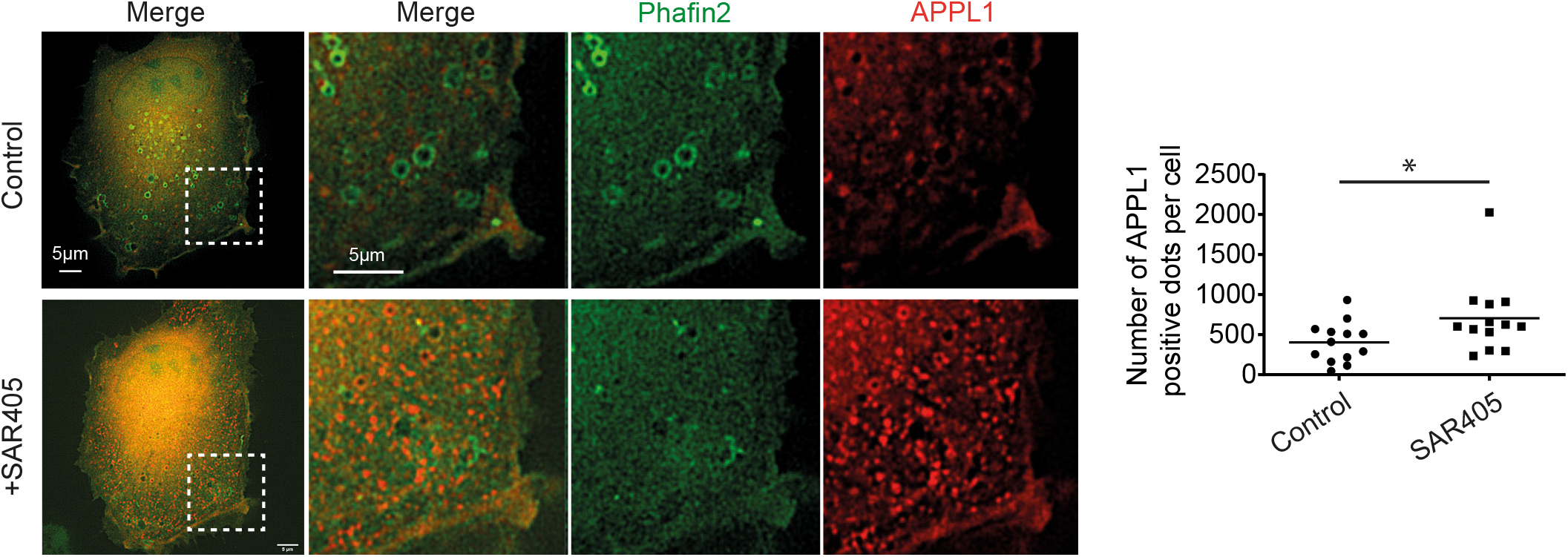
Reconversion of endosomes to the APPL1-positive stage. We performed live cell imaging of Phafin2-GFP and mCherry-APPL1 expressing cells. The number of mCherry-APPL1 positive structures was quantified in the last frame before and in the first frame after addition of the inhibitor. Comparing the number of APPL1-positives structures between the untreated and treated frames, we observed a significant increase of APPL1-positive structures in SAR405 treated conditions (n=13 cells for each condition/ 1 day, Shapiro-Wilk: Control p=0,7605; SAR405 p=0,0031; Mann-Whitney Test p=0,0256).

Movie S1

Live imaging of hTert-RPE1 cells stably expressing mCherry-MyrPalm (magenta) and mNeonGreen-2xFYVE (green) in the presence of VPS34-inhibitor SAR405 (3 μM). The movie shows the formation of a macropinosome as marked by an arrow and follows its progress after formation (scalebar 2 μm).

Movie S2

Live imaging with an oblique plane microscope of HT1080 cells expressing Phafin2-GFP in the presence of SAR405 (3 μM). The movie shows macropinosome formation dynamics in 3D and shows a macropinosome re-fusing with the plasma membrane.

Movie S3

Live imaging of hTert-RPE1 cells stably expressing Phafin2-GFP (green) co-transfected with mCherry-RAB5Q79L (magenta) in the presence of SAR405 (3 μM). The movie shows the recruitment of mCherry-RAB5Q79L to nascent macropinosomes before they re-fuse with the plasma membrane (scalebar 2 μm).

Movie S4

Live imaging of hTert-RPE1 cells co-transfected with mNeonGreen-RAB5a (green) and mCherry-RAB8A (magenta). The movie shows to macropinosomes marked by arrows that transition from mCherry-RAB8A to mNeonGreen-RAB5A (scalebar 2 μm)

Movie S5

Live imaging of hTert-RPE1 cells co-transfected with GFP-RAB5Q79L (green) and mCherry-RAB8A (magenta) in the presence of SAR405 (3 μM). The movie shows nascent macropinosomes positive for both GFP-RAB5Q79L and mCherry-RAB8A (scalebar 2 μm).

Movie S6

Live imaging of hTert-RPE1 cells stably expressing Phafin2-GFP (green) and mCherry-RAB8T22N (magenta) in the presence of SAR405 (3 μM). The movie shows nascent macropinosomes that are unable to re-fuse with the plasma membrane due to inactivation of RAB8a. A macropinosome that forms and undergoes several Phafin2-GFP recruitments is marked with a white arrow (scalebar 2 μm).

## Author contributions

H.Sp. designed experiments, generated constructs, lentivirus and stable cell lines, performed live cell imaging, flow cytometry assays, analyzed data and wrote the original draft. M.S. performed live cell imaging and data analysis. M.M.T. generated constructs and stable cell lines and performed live cell imaging. C.VM. and Y.Y.C. performed exploratory proteomics. K.O.S. performed live cell imaging, generated constructs and wrote image and data processing software. H.St. provided funding, co-supervised the study, and discussed data. K.O.S. and C.R. conceived and supervised the study and contributed to data validation and visualization. All co-authors reviewed and edited the manuscript.

## Acknowledgements

We thank Ling Wang for the qPCR analysis of RAB10, we thank Simona Migliano for the help with performing an APEX screen and proteomics analysis. We thank Jost Enninga for discussions and help during exploratory proteomics. We thank Ulrikke Dahl-Brinch for help with cloning of constructs. We thank Kia Wee Tan for discussions on the project.

K.O.S was supported by a Career grant from the South-Eastern Norway Regional Health Authority (2020038) and a Research Grant from the Research Council of Norway (315103). H.Sp. was supported by an Open Project Grant from the South-Eastern Norway Regional Health Authority (2018081), a Research Grant from the Norwegian Cancer Society (182698) and an Advanced Grant from the European Research Council (788954). This work was partly supported by the Research Council of Norway through its Centres of Excellence funding scheme, project number 179571. C.R. was supported by a Research Grant from the Norwegian Cancer Society (198140).

## Notes

### Competing Interest Statement

The authors have declared no competing interest.

